# Parkinson’s Disease affects the contextual control, but not the expression, of a rapid visuomotor response that initiates visually-guided reaching: Evidence for multiple, interacting motor pathways and implications for motor symptoms in Parkinson’s Disease

**DOI:** 10.1101/2024.11.27.625399

**Authors:** Madeline Gilchrist, Rebecca A. Kozak, Margaret Prenger, Mimma Anello, Kathryne Van Hedger, Penny A MacDonald, Brian D Corneil

## Abstract

Despite significant deficits in voluntary motor control, patients with Parkinson’s disease (PD) can generate reflexive or stimulus-driven movements. How are such spared capabilities realized? Here, we recorded upper limb muscle activity in patients with PD and age-matched healthy controls (HCs) as they reached either toward or away from a visual stimulus. The task promoted express visuomotor responses (EVRs), which are brief bursts of muscle recruitment time-locked (<100 ms) to stimulus presentation that are thought to originate from the midbrain superior colliculus. Across two experiments, we observed a remarkable sparing of the latency and magnitude of EVRs in patients with PD, but a decreased ability for patients with PD to contextually modulate the EVR depending on trial instruction. EVR Magnitudes were strikingly strongly correlated with PD Reaction times and Error rates, despite compromised levels of electromyographic (EMG) recruitment in subsequent phases of muscle activity, which predicted lower Peak velocities. Our results are consistent with a differential influence of PD on parallel-but-interacting subcortical and cortical pathways that converge onto brainstem and spinal circuits during reaching. This differential influence is discriminable even within a single trial in the selective sparing of stimulus-aligned but not movement-aligned muscle recruitment, and has implications for our understanding of the motor and cognitive deficits seen in PD.

## Introduction

Parkinson’s disease (PD) is a multisystem neurodegenerative disorder. Classic motor symptoms of PD, including muscle rigidity, rest tremor, slowed (i.e., bradykinesia) and small amplitude (i.e., hypokinesia) movements are ascribed to degeneration of dopamine-producing neurons in the substantia nigra pars compacta (SNc) and consequent dopamine depletion in caudal motor portions of the dorsal striatum (Fearnley and Lees, 1991). Despite these motor deficits, patients with PD can occasionally still move as quickly as healthy people under certain conditions, referred to as *kinesis paradoxica (KP).* KP is seen under threatening situations and in response to certain, sudden external stimuli, such as in hitting an approaching tennis ball (Glickstein and Stein, 1991; Duysens and Nonnekes, 2021). In the laboratory, seemingly parallel phenomena have been observed when PD patients and healthy participants are equally capable of adjusting reaching trajectories to a visual target that is abruptly displaced just before or soon after movement onset (Desmurget et al., 2004; Merritt et al., 2017). Similarly, despite deficits in voluntary eye movements, PD patients can also generate various reflexive or visually-guided eye movements, such as express saccades, which are the shortest latency, visually-guided saccades normally (Vidailhet et al., 1994; Briand et al., 2001; Cubizolle et al., 2014; Antoniades et al., 2015; Fooken et al., 2022; Riek et al., 2023).

The case of express saccades is particularly interesting, as express saccades require integrity of the midbrain superior colliculus (Schiller et al., 1987; Edelman and Keller, 1996; Dorris et al., 1997; Sparks et al., 2000). The SC also contributes to head and upper limb movements via rich tecto-reticulo-spinal projections (Gandhi and Katnani, 2011; Corneil and Munoz, 2014; Cooper and McPeek, 2021), and this structure has been implicated in the phenomenon of *express visuomotor responses* (EVRs) on neck and upper limb muscles (Corneil et al., 2004; Pruszynski et al., 2010). EVRs are extremely short-latency (100 ms or less) bursts of muscle recruitment that can be measured with electromyography (EMG) and share many response properties with express saccades. Like express saccades, EVRs are tightly locked to the time and location of stimulus onset. EVRs can only be directed *toward* a stimulus, even when subsequent phases of reaching are withheld (Corneil et al., 2008; Wood et al., 2015; Atsma et al., 2018) or proceed in the opposite direction (Chapman and Corneil, 2011; Gu et al., 2016; Contemori et al., 2022). EVRs, like express saccades, are potentiated by high-contrast, low-spatial frequency stimuli (Marino et al., 2015; Wood et al., 2015; Kozak et al., 2019), and various top-down cueing and timing signals (Paré and Munoz, 1996; Schiller et al., 2004; Contemori et al., 2021a, 2021b, 2022). If EVRs are indeed mediated by the SC, then like express saccades they might be spared in PD. We speculated that spared EVRs could underlie the preservation of rapid, reflexive reaching movements in PD. Furthermore, we wondered if PD would have different or similar effects on EVR modulation, and subsequent EMG activity related to potentially-altered kinematics of the executed reach. Most motor symptoms of PD remain poorly understood. Understanding these effects and interactions might clarify some symptoms of PD.

Across two experiments, we measured reach kinematics and recorded upper limb muscle activity with surface EMG in PD patients and age-matched healthy controls (HCs) during a task that required participants to either reach toward (termed a Pro-reach trial) or away from (termed an Anti-reach trial) an emerging visual stimulus (**Fig. 1**). We increased task difficulty in the second experiment by shortening the instruction to prepare for a Pro- or Anti-reach to as little as 500 ms. Anti-reaches dissociate bottom-up, stimulus-mediated EVRs from top-down, goal-directed movement selection and enactment — phases that might relate to later muscle activity. Though EVRs persist in both Pro- and Anti-Reach settings, they are dampened by the Anti-reach instruction (Gu et al., 2016; Kozak et al., 2020), providing evidence of contextual control of the EVR and an opportunity to investigate these interactions in PD patients. Here, we investigate the effect of PD on the expression and contextual control of EVRs, and relate both EVRs and subsequent phases of goal-driven muscle recruitment to the kinematics of the reach itself. We find that patients with PD had a greater speed-accuracy tradeoff than HCs, evidenced by comparable reaction times (RTs) but more frequent reach directional errors, and also had lower Peak reach velocities and longer Movement durations, which seemed consistent with bradykinesia. These kinematic measures were remarkably strongly correlated with different phases of muscle recruitment. EVRs, which were spared in PD, correlated with RT and Error rates, whereas Peak reach velocities and Movement durations were related to subsequent EMG activity following the EVR. This second burst of EMG activity was systematically lower in PD patients. Our results provide insights into the pathophysiology of reaching deficits in PD and motor symptoms more broadly. Whereas PD affects descending motor pathways that depend on nigrostriatal/corticostriatal inputs, it selectively spares tecto-reticulo-spinal pathways that generate reflexive actions to visual stimuli.

**Figure 1.**
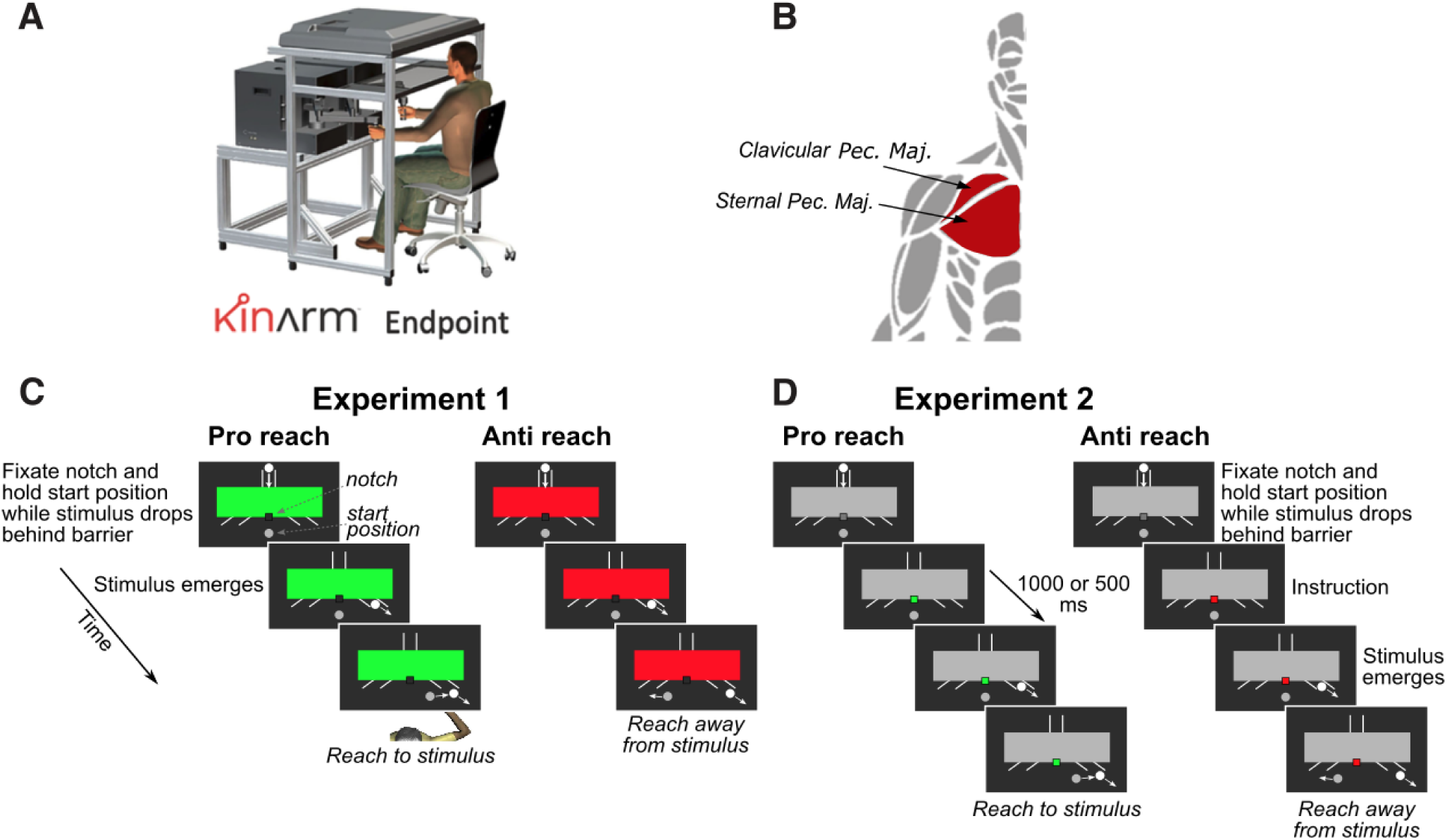
Behavioral paradigms. **A**. Participants interacted with a Kinarm End-point robot, moving a manipulandum with their right hand. Surface electromyographic (EMG) recordings were acquired from the clavicular and sternal heads of right pectoralis major (**B**), which is recruited for leftward movements. **C**. In Experiment 1, we interleaved pro- and anti-reach variants of the emerging target paradigm. At the start of each trial, participants acquired the central start position with their hand (grey circle), and fixated a small notch at the bottom of the barrier. The barrier color presented at the start of each trial conveyed the instruction to reach toward (green barrier, a pro-reach) or away from (red barrier, an anti-reach) the stimulus (white circle) upon its emergence below the barrier in the right or left outlet. This instruction was available to the participant for at least 1500 ms before stimulus emergence. The side of stimulus emergence was randomized, but only right emerging stimuli are depicted here. **D**. In Experiment 2, the sequence was largely the same, except that the instruction was conveyed by the color of a square at the notch (green for pro-reach, red for anti-reach) presented either 1000 or 500 ms before stimulus emergence. Doing so increased task difficulty by reducing instruction time.

## Methods

### Participant demographics

Twenty-six unique participants (12 females, 14 males; Age range: 59-79) who had been diagnosed with idiopathic PD by a movement disorders neurologist were enrolled in this study (**Table 1**). Twenty-five (12 females, 13 males; Age range: 57-79) unique age-matched HCs were also recruited. Of the patients with PD, 16 (8 females, 8 males) completed Experiment 1 and 17 (8 females, 9 males) completed Experiment 2. Four female and 3 male PD patients completed both experiments with at least 12 months between experiments. Eighteen HCs (9 females, 9 males) completed Experiment 1, and 18 HCs (9 females, 9 males) completed Experiment 2. Six female and 5 male HCs completed both experiments with at least 12 months between experiments. HCs were age- and education-matched to within five years of the matched PD patient. Participants were recruited through the Movement Disorders Database at London Health Sciences Center. Exclusion criteria for our PD group included a diagnosis of any neurological disorder other than PD, a history of alcohol or drug abuse, dementia, hallucinations, uncorrected visual deficits including colour blindness, a history of deep brain stimulation treatment, injuries or conditions preventing normal movement of the right arm, and psychiatric disorders, save for mild-moderate depression or anxiety. Exclusion criteria for our HC group were identical to those applied to our PD group save for the fact that none of our HCs had PD. Participants taking cognitive-enhancing medications including donepezil, galantamine, rivastigmine, memantine, or methylphenidate were excluded from participating. Participants were naïve to the purpose of the experiment. All participants provided written informed consent according to the Declaration of Helsinki (1991). All procedures were approved by the Health Sciences Research Ethics Board of the University of Western Ontario (London, Ontario, Canada).

**Table 1:** Demographics, clinical, and questionnaire measures for HCs and PD patients

|  | Experiment 1 |  | Experiment 2 |  |
| --- | --- | --- | --- | --- |
|  | HC<br>(n = 18) | PD<br>(n = 16) | HC<br>(n = 18) | PD<br>(n = 17) |
| Sex = F | 9 (50.00%) | 8 (50.00%) | 9 (50.00%) | 8 (47.06%) |
| Age | 68.39 (5.89) | 68.62 (5.70) | 67.89 (6.01) | 68.18 (5.49) |
| Education (years) | 15.83 (1.72) | 15.75 (1.95) | 15.00 (1.37) | 14.35 (2.83) |
| AmNART | 122.70 (7.62) | 123.76 (8.45) | 124.39 (6.61) | 123.20 (8.52) |
| MoCA | 28.17 (1.69) | 27.88 (1.41) | 28.38 (1.59) | 26.82 (2.13) |
| BDI-II | 1.89 (2.19) | **10.50 (7.71)\*\*** | 2.19 (2.10) | **12.12 (6.67)\*\*** |
| BAI | 0.50 (1.04) | **7.63 (5.14)\*\*\*** | 1.56 (1.63) | **12.35 (10.26)\*\*** |
| BMI | 27.72 (4.61) | 26.62 (4.14) | 28.38 (1.59) | 28.76 (4.24) |
| Handedness = R | 15 (88.24%) | 14 (87.50%) | 15 (93.75%) | 14 (82.35%) |
| PD dominant side = R | -- | 8 (50.00%) | -- | 10 (58.82%) |
| Disease Duration (yr) | -- | 3.80 (4.86) | -- | 5.87 (4.62) |
| LEDD (mg) | -- | 479.69 (354.17) | -- | 633.41 (332.13) |
| N-FOG | -- | 3.75 (6.45) | -- | 5.18 (7.75) |
| UPDRS OFF | 2.88 (3.34) | **31.69 (9.79)\*\*\*** | 2.15 (2.03) | **28.53 (13.55)\*\*\*** |
| UPDRS ON | 2.93 (3.25) | **23.67 (9.42)\*\*\*** | 2.08 (2.06) | **24.03 (14.86)\*\*\*** |

**Note:** Data are presented as mean values with standard deviations in parentheses, except where data are presented in raw values with percentages in parentheses. AmNART = American National Adult Reading Test; MoCA = Montreal Cognitive Assessment; BDI-II = Beck Depression Inventory II; BAI = Beck Anxiety Inventory; BMI = Body Mass Index; LEDD = Levodopa Equivalent Daily Dose; N-FOG = New Freezing of Gait Questionnaire; UPDRS = motor subscale of the Unified Parkinson's Disease Rating Scale; \* $p < .05$ , \*\* $p < .01$ , \*\*\* $p < .001$ .

### Manipulation of dopamine status

All participants were enrolled in a study requiring two visits on separate days, one day off medication (i.e., the OFF session) and one day on medication (i.e., the ON session), following procedures outlined by (Jost et al., 2023). The current manuscript reports data obtained from the OFF session only. The OFF-ON order was counterbalanced across participants. For their OFF session, PD participants were instructed to abstain from all dopaminergic medications including dopamine precursors such as levodopa, aromatic L-amino-acid decarboxylase inhibitors such as carbidopa, and catechol-O-methyltransferase (COMT) inhibitors such as entacapone for a minimum of 12 to a maximum of 18 h, as well as dopamine agonists, such as pramipexole (Mirapex), ropinirole (Requip), or pergolide (Permax), amantadine (Symmetrel), rasagiline (Azilect), and selegiline (Eldepryl or Deprenyl) for 16 to 20 h before testing. For the ON session, PD patients were instructed to take their regular medication as prescribed. HCs were administered a capsule orally, 45 minutes before experimental testing. This capsule was single-blinded, and consisted of either a cornstarch placebo (i.e., the OFF session), or a levocarb capsule consisting of 100 mg levodopa and 25 mg carbidopa (i.e., the ON session). A future manuscript will report on the influence of exogenous dopamine on behaviour and upper limb muscle recruitment.

We obtained a variety of motor and affective measures and briefly mention them here to confirm the effect of dopaminergic modulation, and to better contextualize and compare PD patients and HCs in the OFF state. At the time of recruitment, all participants with PD were taking dopaminergic medication in dosages and manners prescribed by their treating movement disorder neurologists. Levodopa Equivalent Daily Dose (Experiment 1: *M* = 479.69 mg, *SD* = 354.17; Experiment 2: *M* = 633.41 mg, *SD* = 332.13) was calculated using the formula described by Merritt et al. (2017): levodopa dose + levodopa × 1/3 if on entacapone + bromocriptine (mg) × 10 + cabergoline or pramipexole (mg) × 67 + ropinirole (mg) × 20 + pergolide (mg) × 100 + apomorphine (mg) × 8. No HCs were taking dopaminergic therapy. The motor subscale of the Movement Disorder Society (MDS)-Unified Parkinson’s Disease Rating Scale (UPDRS) III was scored by a licensed neurologist with subspecialty training in movement disorders (P.A.M.) to assess the presence and severity of motor symptoms for all patients both off and on dopaminergic medication. Patients with PD had significantly higher MDS-UPDRS III scores in the OFF state compared to the ON state (Experiment 1: *t*(11) = -5.310, *p* < 0.001; Experiment 2: *t*(16) = -5.427, *p* < 0.001). HCs were also screened to rule out undiagnosed neurological illness using the MDS-UPDRS III, also evaluated by a movement disorder specialist. There was no difference in the MDS-UPDRS III scores of the HC participants between the OFF and ON states (Experiment 1: *t*(13) = -1.883, *p* = 0.082; Experiment 2: *t*(12) = -0.201, *p* = 0.844). Affective measures were obtained for PD patients and HCs (**Table 1**). T-tests revealed PD patients had significantly higher Beck Depression Inventory-II (BDI-II) and Beck Anxiety Inventory (BAI) scores than HCs, which was expected given that mild-moderate anxiety and depression are symptoms of PD (Broen et al., 2016).

### Apparatus and behavioural tasks

Participants were seated in a KINARM End-Point Lab (BKIN Technologies, Kingston, ON Canada), and performed visually-guided reaching movements with their right arm as they gripped a manipulandum (**Fig. 1A**). Visual stimuli were projected onto an upward facing mirror from a downward facing monitor in a custom built-in projector (PROPixx projector by VPixx, Saint-Bruno, QC, Canada). A shield below the mirror occluded direct vision of the hand, but real-time hand position was represented on the monitor via a real-time cursor projected onto the screen. Surface EMG electrodes (Bagnoli-8 system, Delsys Inc., Boston MA) were placed over the clavicular and sternal heads of the right pectoralis major (**Fig. 1B**) — a muscle that is recruited for cross-body (i.e., leftward, in our case) planar reaching. During both experiments, we applied a constant load of 5 N to the right and 2 N toward the participant to increase the baseline recruitment of the muscle of interest. Because of this baseline recruitment, the EVR consists of either an increase or decrease in muscle recruitment following stimulus presentation in, or opposite to, the muscle’s preferred direction of movement respectively. EVRs are tuned in the same manner and direction as movement-aligned activity (Pruszynski et al., 2010; Gu et al., 2019; Selen et al., 2023). This constant load is relatively small and was well tolerated. For comparison, gravity exerts a force of 29.4 N on an outstretched arm with a mass of 3 kg. Another benefit of the constant load is that it reduced the resting tremor in PD patients, making it easier to identify movement onset.

Both Experiment 1 (**Fig. 1C**) and Experiment 2 (**Fig. 1D**) were constructed as variants of the Emerging Target Paradigm (Kozak et al., 2020). In this paradigm, a stimulus drops down behind a barrier, and subsequently emerges in motion at the right or left outlet; participants reach in response to stimulus emergence. Rapid responses on the upper arm muscle in the direction of the emerging stimulus, or EVRs, are readily generated in this paradigm in the majority of participants studied to date (Contemori et al., 2021a, 2021b, 2022; Kozak and Corneil, 2021; Kearsley et al., 2022). Because the arm is stable at the time of stimulus emergence, muscle recruitment arising from stimulus emergence is more easily distinguished than if the arm was in motion, as in an online reach correction paradigm. Both experiments intermixed the instruction to reach toward (i.e., a Pro-reach) or away from (i.e., an Anti-reach) the emerging stimulus. In Experiment 2, we also manipulated the time that this instruction was available prior to the emergence of the stimulus to investigate the effect of decreased instruction time. For brevity, we will describe the paradigms used in Experiments 1 and 2 together.

Trials in both experiments began with a stimulus that appeared in a vertical channel above a barrier (**Fig. 1C, D**). Participants were required to move the real-time cursor to a central start location. Participants were instructed at the start of each trial to look at a small notch (Experiment 1) or small gray square (Experiment 2) located at the bottom of the barrier. The stimulus then dropped down the vertical channel, disappeared behind the barrier for 1000 ms before emerging from below the occluder in continuous oblique-downward motion from either the right or left outlet. The outlets were approximately 20 cm lateral to and slightly above the central start location. In Experiment 1, the instruction for how to respond to stimulus emergence was conveyed by the colour of the barrier that was present for the entire trial, beginning at least 1500 ms before stimulus emergence, with green indicating Pro-reach and red signalling Anti-reach trials. For brevity, we refer to the instruction time in Experiment 1 as being 1500 ms in the Results. In Experiment 2, the instruction for how to respond to the emerging stimulus was conveyed by the colour of the small fixation square, which appeared in the centre of the screen, was initially gray and, either 1000 ms or 500 ms prior to the emergence of the stimulus, changed to green or red. On Pro-reach trials, participants were instructed to reach toward and intercept the emerging stimulus. On Anti-reach trials, participants were instructed to reach away from the stimulus. At the end of each trial, the following feedback was written across the occluder in the inter-trial interval: a) “HIT” for an intercepted stimulus on Pro-reach trials or for reaches that went at least 10 cm away from the emerging stimulus on Anti-reach trials, b) “WRONG WAY” for a movement that went in the wrong direction and was not corrected, or c) “MISS” for a reach that went in the correct direction but did not intercept the stimulus on Pro-reach trials, or did not achieve the required 10 cm movement on Anti-reach trials. In Experiment 1, participants completed 5 blocks of 100 trials each, for 500 trials total. In Experiment 2, participants completed 5 blocks of 104 trials each, for 520 trials total. Within each block, all trial combinations, a) Pro- or Anti-reach, b) stimulus left or right, and, for Experiment 2, c) 1000 or 500 ms of instruction time were pseudorandomly varied to be presented an equal number of times. In total, we had 125 repeats of each unique trial condition in Experiment 1, and 65 repeats of each unique trial condition in Experiment 2.

### Data acquisition and analysis

Data acquisition techniques resemble those published previously (Kozak and Corneil, 2021; Kearsley et al., 2022). Kinematic data were sampled at 1 kHz by the KINARM platform. The precise time of stimulus emergence below the barrier was indicated by the presentation of an additional visual stimulus which was unseen by the participant. A photodiode was placed over the location of this additional visual stimulus, and all kinematic and EMG data were aligned to photodiode onset. Kinematic data from each trial were examined via customized MATLAB GUIs that permitted the exclusion of clearly atypical trials (e.g., when the participant failed to respond, moved the arm well before stimulus emergence, made multi-step reaches in a given direction, or reached in the wrong direction without any subsequent corrective movement). For each trial, we derived a number of reach metrics: movement RT, movement trajectory and endpoint, tangential Peak velocity, and Movement duration. RT was calculated as the time from stimulus emergence to reach initiation, defined as the time at which tangential hand velocity exceeded 5% of the Peak velocity on that trial. The end of the reach was defined as the point at which tangential hand velocity reached zero. Peak velocity and Movement duration were determined for the movement segment between RT and movement offset.

A customized MATLAB GUI allowed us to demarcate incorrect movements in the wrong direction for each trial, even if the peak velocity during small incorrect movements did not exceed 5%. All participants commonly made small reaching errors on Anti-reach trials, moving sometimes quite subtly toward the emerging stimulus before correctly reaching in the opposite direction (**Fig. 2**). Sequences where the participant initially moved in the wrong direction and then subsequently reached in the correct direction were retained for further analysis of error magnitude. Given that the reaching movements of PD patients are often of lower amplitude or hypokinetic, we extracted the proportional amplitude of these erroneous reaching movements relative to each participant’s average reach amplitude on Pro-reach trials in that direction. Thus, a “10% error” is a reaching error where the participant moved 10% of the way toward the emerging stimulus, relative to their average movement amplitude on Pro-reach trials. For the kinematic results presented in this manuscript, we reclassified “<5% error” Anti-reach trials as correctly-performed trials as long as the participant made a corrective reach in the correct direction which crossed the centre within 500 ms of stimulus onset, and for such trials, we extracted movement parameters (i.e., RT, *C*/*d’,* Peak velocity, and Movement duration) for the reach component in which participants were correctly moving away from the emerging stimulus.

**Figure 2.**
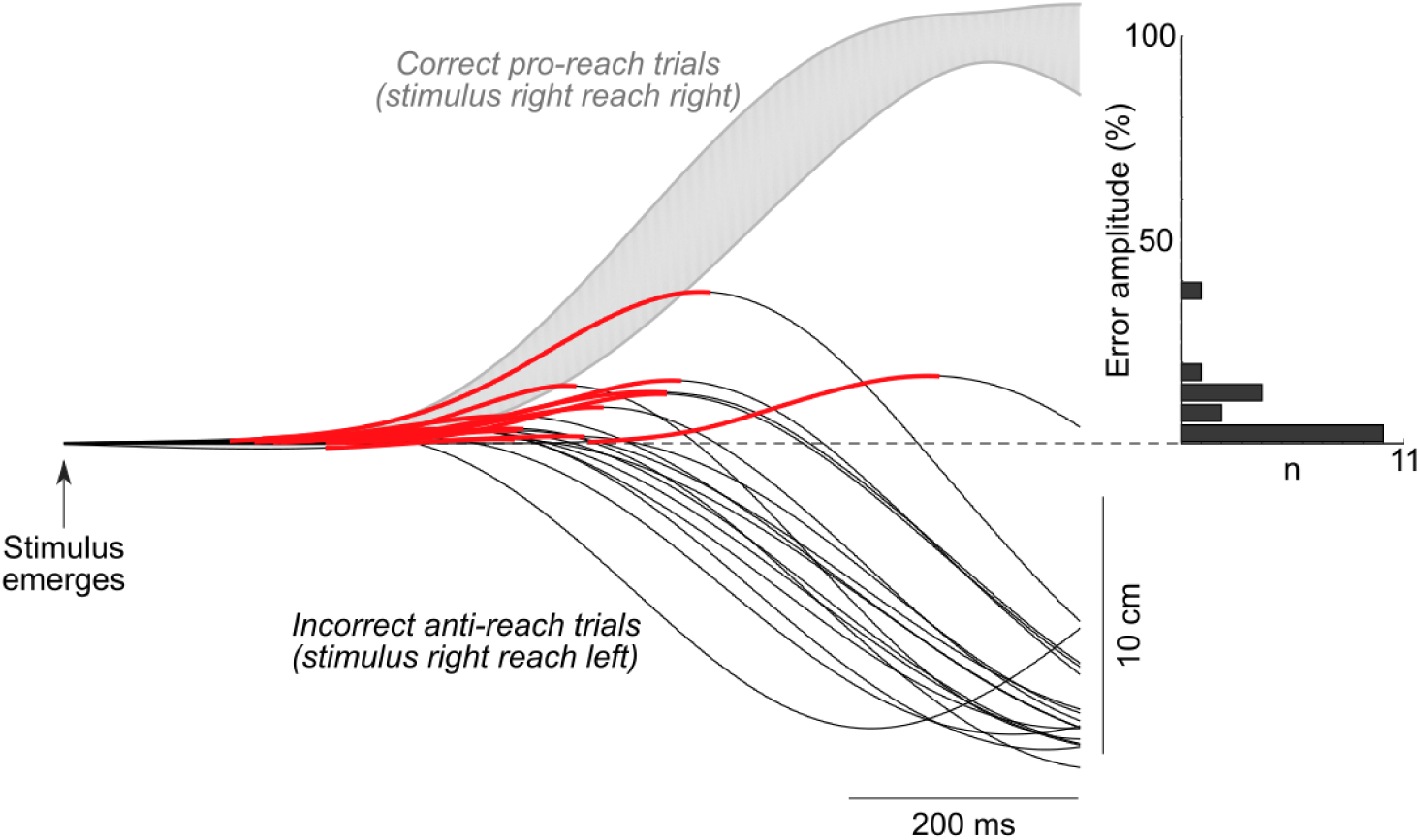
Calculation of anti-reach error amplitude, showing data from a single PD patient in Experiment 2 with 500 ms of instruction time. Thin traces depict horizontal hand position through time for individual anti-reach trials where the patient moved slightly toward a stimulus that emerged to the right before correctly moving to the left (right = up); the incorrect movement component is coloured in red. Grey contour depicts the average hand position through time for right pro-reach trials, subtended by the standard deviation. Error amplitudes on anti-reach trials were scaled to the average reach amplitude on pro-reach trials. The histogram shows that most anti-reach errors for this patient moved < 25% of the way to the stimulus before being corrected.

Individual trials were excluded if the RT was shorter than 130 ms (indicating anticipatory movement). This cutoff was supported by an analysis showing that Pro-reach trials with RTs less than 130 ms were correct 50% of the time, whereas those with RTs greater than 130 ms were correct >80% of the time. We also excluded trials with RTs greater than 500 ms, as these were rare and indicated distraction or a lack of vigilance. Across both experiments, 12% and 6% of trials were excluded for PD patients and HCs, respectively.

After trial exclusion, for both Pro-reach and Anti-reach trials, we calculated Error rate, as well as the signal detection parameters response Criterion (*C*) and response Discrimination (*d’*). *C* and *d’* take into account both the hit rate (i.e., the proportion of correct reaches on Pro-reach trials) and the false alarm rate (i.e., the rate of erroneous reaches toward the stimulus on Anti-reach trials) providing a measure of response criterion and discrimination of Pro- and Anti-reach cues reflected in the behavioural response. For *C*, higher values suggest a more conservative internal threshold or bias used to determine whether a response signal (i.e., Pro-reach) is present among noise (i.e. Anti-reach) signals, resulting in less of a likelihood that a participant would reach towards the stimulus on Anti-reach trials, at the cost of also missing more Pro-reach trials. For *d’*, higher values indicate greater ability to modulate the bottom-up, habitual movement toward the emerging visual stimulus by the top-down instruction, which results in better sensitivity of their responses. Conversely, a lower value implies reduced ability to modulate behaviour according to instructions. Response regulation ability measured with *C* and *d’* should be considered in the context of RT data, however, to assess for speed-accuracy trade offs in this task. The latter also provides information about a participant’s ability to modulate bottom-up influences on behaviour related to reach instructions.

EMG data from the electrode on the sternal head of the pectoralis major muscle were used for analysis unless there was a disruption to data collection (e.g., the electrode came loose during the study), in which case the recordings from the clavicular head of the pectoralis major muscle were used, instead. EMG data were amplified by 1000 and sampled at 1 kHz by the Bagnoli 8-system, which has a bandpass filter between 20 to 450 Hz. Offline, EMG signals were full-wave rectified, and smoothed with a 7-point smoothing function. For the analysis of the EVR, trials were considered correct as long as either a) reaches were oriented to the correct side within 500 ms of stimulus onset or b) participants initially moved in the incorrect direction but made a corrective reach in the appropriate direction which passed the centre position within 500 ms of stimulus onset. This is a less stringent inclusion criteria than described above for the kinematic analyses, given that the aim of the EMG analyses is to investigate the modulation of the EVR according to task instruction, and therefore all trials on which instructions were clearly understood and the more automatic reach towards the stimulus was overcome before our RT cutoff (i.e., 500 ms) were of interest.

When possible, the magnitude of EMG recruitment is reported in source microvolts. Comparisons of the magnitude of recruitment for PD patients versus HCs required that we first normalize EMG data to the level of recruitment in the 500 ms preceding the stimulus emergence, when participants were holding their arms stable against the background load. The EVR is the first burst of muscle activity that appears between 80 and 120 ms after stimulus appearance. The EVR is directionally-tuned to the stimulus, and therefore appears as increased activity on the right pectoralis major when the stimulus appears on the left, and decreased activity when the stimulus appears on the right, relative to baseline activation. Detection of EVRs follow previously established methods (Corneil et al., 2004; Pruszynski et al., 2010) relying on a time-series receiver operating characteristic (ROC) analysis. Separate time-series ROCs were run on data for each task variant (e.g., separate time-series ROCs were run on pro- or anti-reaches in Experiment 1, and on pro- and anti-reaches in Experiment 2 for both the 1000 ms and 500 ms instruction time). Briefly, at each timepoint, from 100 ms before, to 200 ms after stimulus emergence, this method compares the distribution of muscle recruitment across all trials where the stimulus emerged to the left or right, and calculates the area-under-curve (AUC) metric; this value indicates the likelihood of discriminating the side of stimulus emergence based on EMG activity alone. A value of 0.5 indicates chance discrimination, whereas values of 1.0 or 0.0 indicate perfectly correct or incorrect discrimination, separately. The discrimination time of a candidate EVR was determined to be the first timepoint in which 8 of the next 10 AUC values exceeded 0.6. If this discrimination time was between 80 and 120 ms after stimulus emergence, then muscle recruitment was deemed to be exhibiting an EVR, and the discrimination time was taken as EVR Latency. EVR Magnitude was taken as the integral over the next 30 ms of the positive difference in mean EMG activity for leftward versus rightward stimulus emergence.

We are also interested in the degree to which instruction to prepare for a Pro-reach vs. Anti-Reach trial influences EVR magnitude. We therefore derived a Modulation index, which was calculated as the difference in EVR Magnitude on Pro-reach vs. Anti-reach trials, divided by the sum of the magnitudes on Pro-reach and Anti-reach trials. The Modulation index is bounded between -1 (EVR on Anti-reach trials but not Pro-reach trials) to 1 (EVR on Pro-reach trials but not on Anti-reach trials); a value of 0 means equal magnitude EVRs are observed on Pro-reach and Anti-reach trials.

Finally, recruitment during the EVR interval (80-120 ms) is followed by subsequent periods of recruitment which persist up to and following movement onset. We therefore also examined the profile of muscle recruitment after the EVR, and examined muscle recruitment after re-aligning all data on movement onset. To obtain a magnitude value for movement-aligned data that is comparable to the stimulus-aligned EVR (i.e., Reach burst magnitude), we first identified the peak of the difference curve of averaged movement-aligned EMG data for leftward and rightward movements in the 100 ms before movement onset, and then calculated the reach burst magnitude as the integral of the difference curve from 15 ms before, to 15 ms after the peak.

### Statistical analysis

Linear mixed models were used to quantify main effects and interactions. Outliers were identified as data points more than 3 *SD* above or below the group mean, and removed from analysis where indicated below. Linear mixed models were chosen over repeated-measures analysis of variance (ANOVA) because of their ability to account for missing data (e.g., if a participant exhibited an EVR in one condition but not another). Unlike ANOVAs, linear mixed models do not use list-wise deletion in the case of missing data points, allowing us to maximize the power and reduce the bias of our analysis. The Satterthwaite method was applied to estimate degrees of freedom and generate *p*-values for the mixed model analyses. We investigated the effect of Group (PD vs. HCs), Trial type (Pro-reach vs. Anti-reach) and Instruction time (1500 ms in Experiment 1 vs. 1000 ms in Experiment 2 vs. 500 ms in Experiment 2), specifying these as fixed effects and participant ID as a random effect in the linear mixed models. Post-hoc orthogonal contrasts with the Bonferroni correction method for multiple comparisons were used to investigate significant interactions between predictor variables. In the event of non-significant Group effects using frequentist linear mixed models, we followed up with Bayesian *t*-tests to evaluate the relative evidence in support of the null hypothesis and alternate hypothesis. Bayesian *t*-tests allow us to make conclusions about a true absence of an effect, which is not possible with frequentist statistics because of the problem of dissimilar error rates of Type I and Type II errors (0.2 for falsely accepting the null and 0.05 for falsely supporting the alternative hypothesis) (Dienes and Mclatchie, 2018; Keysers et al., 2020). A Bayes factor (*BF_10_*) below 3 indicates strong support in favour of the null hypothesis and a Bayes factor above 3 indicates strong support in favour of the alternative hypothesis (Jeffreys, 1998). We investigated the effect of the EVR and Reach-burst magnitude on reach performance, correlating EVR Magnitude and EVR Modulation with RT, Error rate, *C*, *d*’, Peak velocity, and Movement duration. Correlations were completed separately for trials with 1500 ms, 1000 ms, and 500 ms of instruction time. Data processing was done in MATLAB (R2014b), and statistical analyses were performed in Jamovi 2.3.21.0.

## Results

Both PD patients and HCs performed the tasks proficiently, with 88% and 94% of trials performed meeting RT criteria for further analyses respectively. On Anti-reach trials, both PD patients and HCs often moved initially toward the emerging stimulus, as shown in **Fig**. **2**, for a representative PD participant. Such erroneous initial reaches were corrected mid-flight, so that a subsequent reach moved the hand appropriately away from the emerging stimulus. Across both groups and experiments, we quantified the proportional horizontal movement amplitude of erroneous reaches relative to the amplitude of each participant’s average pro-reaches. In both groups, the vast majority of erroneous reaches on Anti-reach trials were corrected before participants reached half of the distance of their average Pro-reach (Experiment 1, 1500 ms: PD = 92.3%, HC = 90.7%; Experiment 2, 1000 ms: PD = 95.1%, HC = 92.7%; Experiment 2, 500 ms: PD = 94.5%, HC = 90.9%). From this, it is clear that both groups consolidated the instruction on Anti-reach trials adequately, consistent with their normal MoCA scores (**Table 1**). Further details about Error rates are provided below.

### RT, Error rate, C, d’

Linear Mixed Models with Group, Trial type, and Instruction time as fixed factors, and Subject ID Intercept as a random factor were performed on RT and Error rate (**Fig. 3A, 3B**). We observed a main effect of Trial type (*β* = -63.69, *SE* = 2.311, *p* < 0.001, *95% CI* [-68.22, -59.16]), as participants had shorter RTs on Pro-reach than Anti-reach trials (*M_Pro_*= 220.1 ms, *SD_Pro_* = 34.98l; *M_Anti_* = 283.8 ms, *SD_Anti_* = 45.47). There was also a significant effect of Instruction time (*β* = -22.65, *SE* = 8.02, *p* = 0.006, *95% CI* [-38.36, -6.927]), as participants responded faster with longer instruction times (*M_1500_* = 237.0 ms, *SD_1500_* = 42.96; *M_1000_* = 241.9 ms, *SD_1000_* = 44.11; *M_500_* = 276.4 ms, *SD_500_* = 57.50). There was a significant interaction between Trial type and Instruction time (*β* = 11.00, *SE* = 4.928, *p* = 0.027, *95% CI* [1.344, 20.66]), and post-hoc comparisons revealed that RTs were significantly shorter with 1500 ms (*p_Pro_* = 0.008; *p_Anti_* < 0.001) and 1000 ms (*p_Pro_* < 0.001; *p_Anti_* < 0.001) compared to 500 ms of instruction time, though the 1500 ms and 1000 ms conditions did not differ from one another (*p_Pro_* = 1.000; *p_Anti_* = 1.000). We found no main effect of Group on RT (*β* = -10.00, *SE* = 8.333, *p* = 0.234, *95% CI* [-26.34, 6.329]). If anything, PD patients tended to have shorter RTs compared to HCs though not significantly (*M_PD_* = 247.0 ms, *SD_PD_* = 48.79; *M_HC_* = 256.6 ms, *SD_HC_*= 53.79). The Bayes factor for the effect of Group indicated some support for the null hypothesis (*BF_10_* = 0.353).

**Figure 3.**
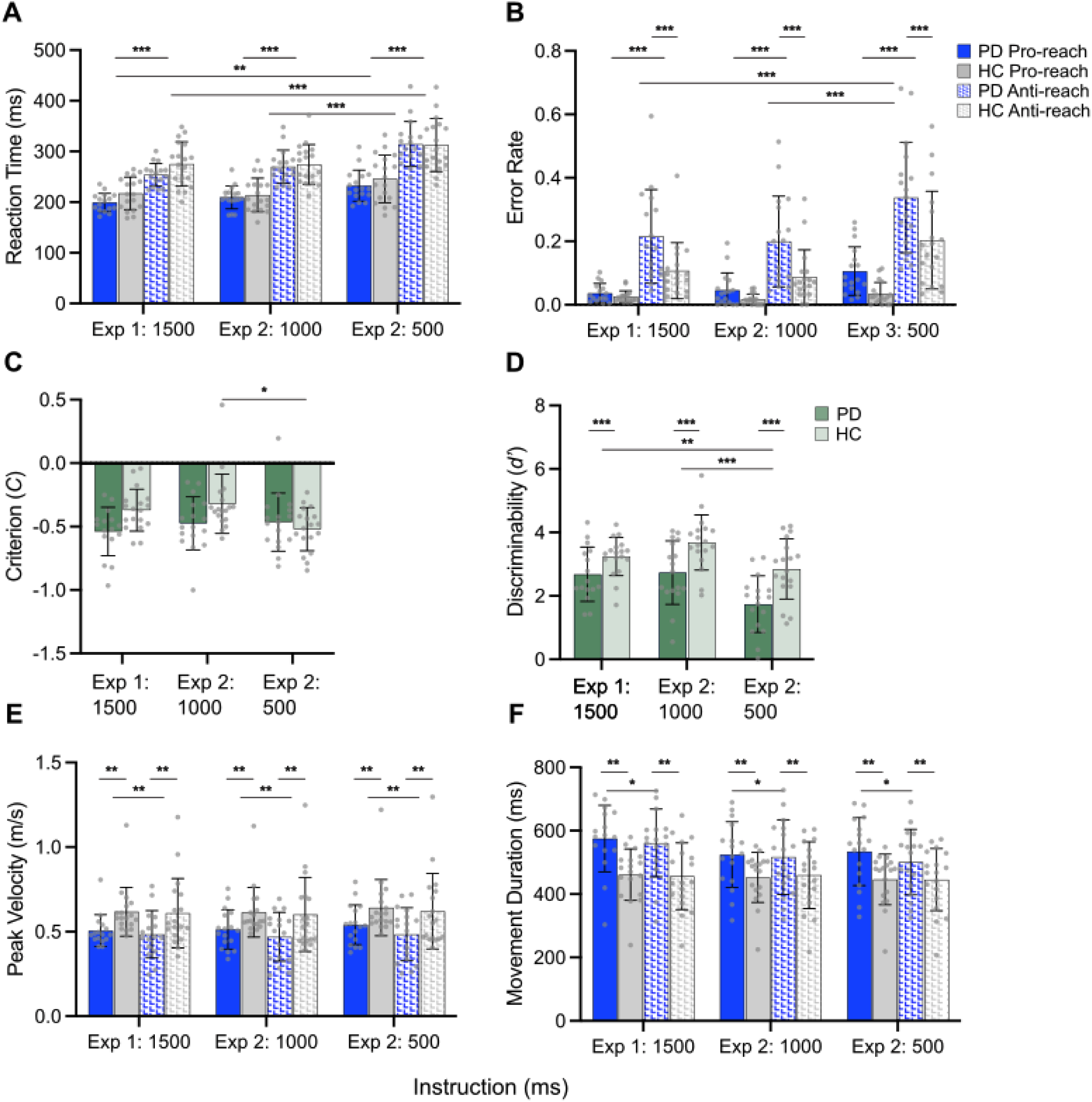
A. Participants had shorter RTs on Pro-reach trials than on Anti-reach trials, and shorter RTs with 1500 ms and 1000 ms of instruction time relative to 500 ms. **B.** PD participants made more errors than HCs on Anti-reach trials. Participants made fewer errors on Pro-reach trials than on Anti-reach trials, and fewer errors on Anti-reach trials with 1500 ms and 1000 ms of instruction time than with 500 ms. **C.** HCs had higher *C* scores with 1000 ms of instruction than with 500 ms of instruction. **D.** PD participants had lower *d’* scores than HCs. Participants had higher *d’* scores with 1500 ms and 1000 ms of instruction time than with 500 ms. **E.** PD participants had lower Peak velocities than HCs. Participants had higher Peak velocity on Pro-reach trials than Anti-reach trials. **F.** PD participants had longer Movement durations than HCs. PD participants had longer Movement durations on Pro-reach trials than Anti-reach trials. Bars represent *M*, error bars represent *SD*, *\*p < 0.05*, *\*\*p < 0.01*, *\*\*\*p < 0.001*.

Next, we examined the rate at which participants reached in the wrong direction (**Fig. 3B**). There was a significant effect of Group on Error rate (*β* = 0.078, *SE* = 0.019, *p* < 0.001, *95% CI* [0.040, 0.115]). PD patients made significantly more directional errors than HCs (*M_PD_* = 0.157, *SD_PD_* = 0.157; *M_HC_* = 0.079, *SD_HC_*= 0.103). As expected, all participants committed fewer reaching errors on Pro-reach than on Anti-reach trials (*β* = -0.148, *SE* = 0.011, *p* < 0.001, *95% CI* [-0.169, -0.127]; *M_Pro_* = 0.044, *SD_Pro_* = 0.052; *M_Anti_*= 0.190, *SD_Anti_*_I_ = 0.156). There was also a significant interaction between Group and Trial type (*β* = -0.081, *SE* = 0.021, *p* < 0.001, *95% CI* [-0.122, -0.039]). Post-hoc comparisons clarified this interaction as resulting from a significant effect of Group on Anti-reach (*p* < 0.001) but not on Pro-reach (*p* = 0.586) trials, with PD patients committing more errors than HCs on Anti-reach but not on Pro-reach trials. There was an effect of Instruction time on Error rate (*β* = 0.079, *SE* = 0.014, *p* < 0.001, *95% CI* [0.052, 0.105]), as participants made fewer directional errors with longer instruction time (*M_1500_* = 0.094, *SD_1500_* = 0.113; *M_1000_* = 0.086, *SD_1000_* = 0.110; *M_500_* = 0.169, *SD_500_* = 0.166). There was also a significant interaction between Trial type and Instruction time (*β* = -0.079, *SE* = 0.022, *p* < 0.001, *95% CI* [-0.123, -0.053]). Post-hoc comparisons indicated that participants made significantly more errors on Anti-reach trials, but not Pro-reach trials, with 1500 ms vs 500 ms (*p_Pro_* = 1.000; *p_Anti_* < 0.001), and 1000 ms vs 500 ms (*p_Pro_* = 0.506; *p_Anti_* < 0.001) of instruction time, but there were no differences between trials with 1500 ms and 1000 ms of instruction time for either Pro-reach or Anti-reach trials (*p_Pro_* = 1.000; *p_Anti_* = 1.000). This finding is consistent with these trials being more challenging due to lesser time for consolidation of the task instruction.

Across both experiments, PD patients revealed a pattern of a) equivalent RTs, but b) more reaching errors, relative to HCs. These results suggested a speed-accuracy trade-off. Accordingly, we analyzed measures of response bias (*C*) and response regulation or discriminability (*d’*) between Pro-reach and Anti-reach trials, relying on concepts from signal detection theory (**Fig. 3C, 3D**). Consistent with speeded responding yet higher Error rate in PD patients, we observed significant effects of Group on *C* (*β* = -0.089, *SE* = 0.041, *p* = 0.035, *95% CI* [-0.169, -0.008]) and *d’* (*β* = -0.874, *SE* = 0.204, *p* < 0.001, *95% CI* [-1.274, -0.474]), reflecting more liberal response criteria and poorer response regulation for PD patients compared to HCs. PD patients responded as quickly as HCs at the expense of poorer response regulation between Pro-reach and Anti-reach trials than HCs, indicated by lower *C* (*M_PD_* = -0.492, *SD_PD_* = 0.209; *M_HC_* = -0.404, *SD_HC_* = 0.207) and *d’* scores (*M_PD_* = 2.381, *SD_PD_* = 1.013; *M_HC_* = 3.261, *SD_HC_*= 0.875), consistent with a higher speed/accuracy trade-off in patients with PD. There was also a main effect of Instruction time on *C* (*β* = -0.040, *SE* = 0.010, *p* < 0.001, *95% CI* [-0.060, -0.021]) and *d’* (*β* = -0.791, *SE* = 0.125, *p* < 0.001, *95% CI* [-1.036, -0.547]), as participants had higher *C* (*M_1500_* = -0.450, *SD_1500_* = 0.194; *M_1000_*= -0.395, *SD_1000_* = 0.232; *M_500_* = -0.493, *SD_500_* = 0.200;) and *d’* (*M_1500_* = 2.981, *SD_1500_* = 0.773; *M_500_* = 2.313, *SD_500_* = 1.069; *M_1000_* = 3.226, *SD_1000_* = 1.040) scores with longer instruction, indicating that response regulation increases with greater time to consolidate the instructional cue. There was a significant interaction effect between Group and Instruction time on *C* (*β* = 0.217, *SE* = 0.080, *p* = 0.008, *95% CI* [0.060, 0.375]). Post hoc comparisons with Bonferroni correction revealed that HCs, but not PD participants had higher *C* scores with 1000 ms than 500 ms of instruction (*p_PD_* = 1.000; *p_HC_* = 0.043), with no significant differences between 1500 ms and 1000 ms (*p_PD_* = 1.000; *p_HC_* = 1.000), or 1500 ms and 500 ms of instruction time (*p_PD_* = 1.000; *p_HC_* = 0.414). The effect of Group on *C* did not survive correction for multiple comparisons with 1500 ms of instruction (*p* = 0.254), 1000 ms of instruction (*p* = 0.383), or 500 ms of instruction (*p* = 1.000), though the Bayes factor (*BF_10_* = 3.092) indicated moderate support for the alternative hypothesis. Post hoc comparisons of *d’* revealed that participants had higher *d’* scores with 1500 ms (*p* = 0.005) and 1000 ms (*p* < 0.001) of instruction compared to 500 ms, but there was no difference between trials with 1500 ms and 1000 ms of instruction time (*p* = 0.691).

### Peak velocity and Movement duration

The analyses of Peak velocity and Movement duration are analogous to those of RT and Error rate. We observed a significant main effect of Group on Peak velocity (**Fig. 3E**; *β* = -0.118, *SE* = 0.038, *p* = 0.003, *95% CI* [-0.193, -0.043]), with PD patients achieving lower velocities than HCs (*M_PD_*= 0.499 m/s, *SD_PD_* = 0.129; *M_HC_* = 0.616 m/s, *SD_HC_* = 0.183). There was a significant main effect of Trial type (*β* = 0.027, *SE* = 0.009, *p* = 0.002, *95% CI* [0.010, 0.044]), with higher velocities on Pro-reach than Anti-reach trials (*M_Pro_ =* 0.573 m/s*, SD_Pro_ =* 0.142; *M_Anti_ =* 0.546 m/s, *SD_Anti_ =* 0.193). The effect of Instruction time on Peak velocity was not significant (*β* = -0.007*, SE* = 0.037*, p* = 0.856*, 95% CI* [-0.078, 0.065]). Participants reached similar velocities with 1500 ms, 1000 ms, and 500 ms of instruction time (*M_1500_* = 0.557 m/s, *SD_1500_* = 0.161; *M_1000_* = 0.550 m/s, *SD_1000_* = 0.170; *M_500_* = 0.572 m/s, *SD_500_*= 0.179). Consistent with findings using Peak velocity as the dependent measure, a significant effect of Group on Movement duration was noted (**Fig. 3F**; *β* = 81.92, *SE* = 24.36, *p* = 0.001, *95% CI* [34.18, 129.67), with longer movement durations in PD patients than HCs (*M_PD_*= 534.9 ms, *SD_PD_* = 107.7; *M_HC_* = 453.6 ms, *SD_HC_* = 90.33). A main effect of Trial type (*β* = 8.921, *SE* = 4.226, *p* = 0.037, *95% CI* [0.638, 17.20), was explained by longer movement durations on pro-reaches than anti-reaches (*M_Pro_ =* 497.0 ms*, SD_Pro_ =* 102.8; *M_Anti_ =*488.4 ms, *SD_Anti_ =* 111.1). We found a significant interaction between Group and Trial type (*β* = 19.00, *SE* = 8.452, *p* = 0.026, *95% CI* [2.432, 35.56]), clarified through Bonferonni-corrected post-hoc comparisons indicating that Movement durations were significantly longer on Pro-reach trials than Anti-reach trials for PD patients (*p* = 0.018) but not HCs (*p* = 1.000). Finally, there was no effect of Instruction time on Movement duration (*β* = 28.51*, SE* = 23.31*, p* = 0.226*, 95% CI* [-17.18, 74.19]), and participants had similar reaching times across all conditions (*M_1500_* = 510.4 ms, *SD_1500_* = 112.1; *M_1000_* = 487.4 ms, *SD_1000_* = 105.1; *M_500_* = 480.8 ms, *SD_500_* = 102.7). Overall, the findings with Peak velocity and Movement duration are consistent with PD-related bradykinesia.

### EVR Prevalence, Latency, and Magnitude

We now turn to the patterns of muscle recruitment during the first wave of activity in response to stimulus emergence. **Figure 4** shows the muscle activity recorded from the right pectoralis major muscle from two PD patients and two HCs, showing activity on all four trial types from Experiment 1. Focusing first on Pro-reach trials (first, second, and fifth columns in **Fig. 4**), the EVR is the first burst of muscle recruitment, consisting of an increase or decrease in activity following stimulus emergence to the left or right respectively. For both the PD and HC participants, such recruitment began in less than 100 ms, reaching magnitudes that neared or exceeded subsequent levels of recruitment. The trial-by-trial timing of such recruitment was more closely aligned to stimulus emergence than reach movement onset, but larger magnitude EVRs preceded shorter-latency reaches. These features are consistent with previous reports (Pruszynski et al., 2010; Wood et al., 2015).

**Figure 4.**
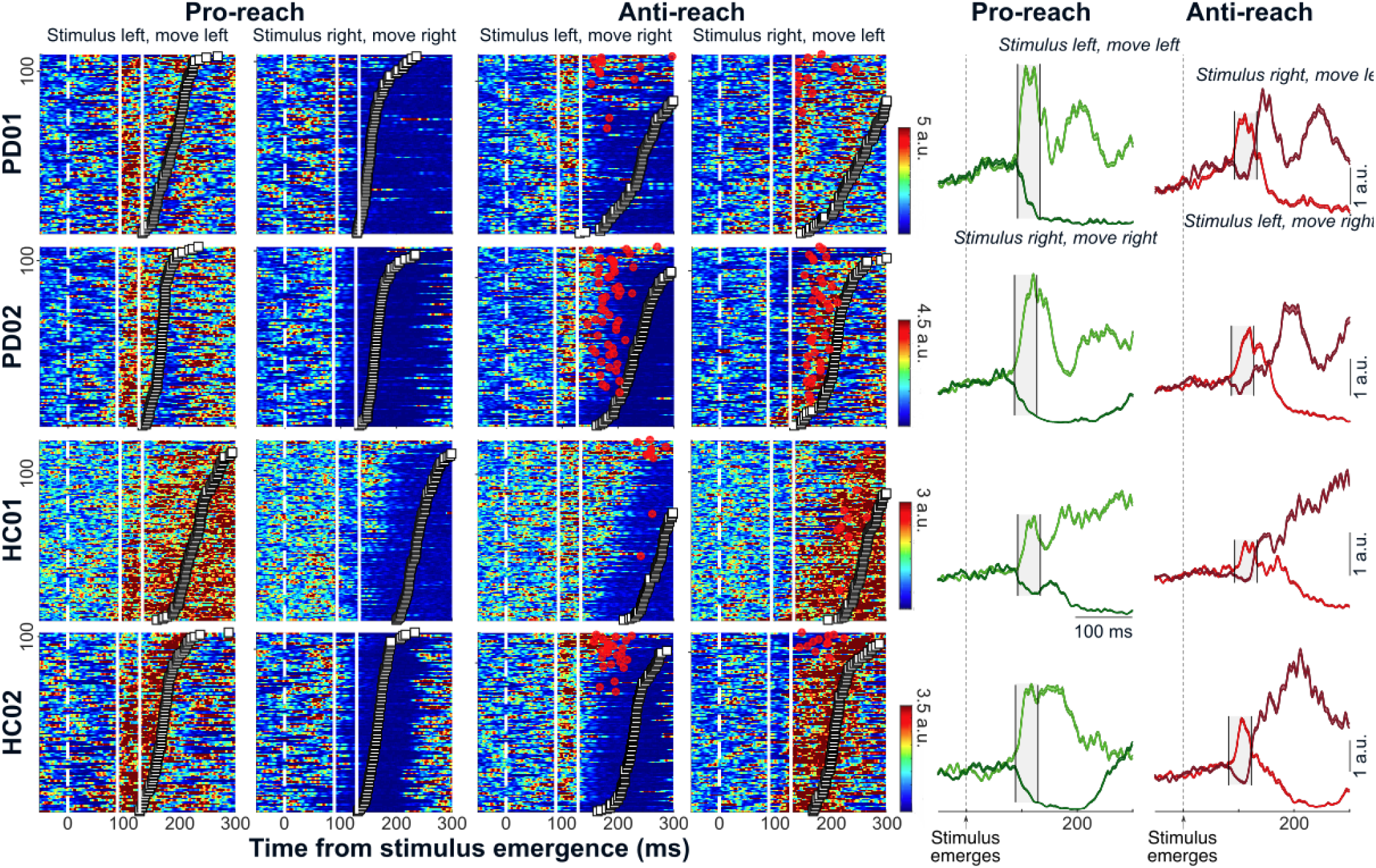
EVRs on the right pectoralis muscle. Example data from Experiment 1 from two PD patients (top two rows) and from two HCs (bottom two rows). From left to right, first two columns show heatmaps of muscle activity for pro-reaches, third and fourth columns show heatmaps of muscle activity for anti-reaches, and fifth and sixth columns depict mean EMG activity subtended by SE for pro- and anti-reaches, respectively. Each heatmap depicts trial-by-trial EMG activity aligned to stimulus emergence (*vertical white dashed lines*), with trials ordered by RT in the correct direction (*white squares*). *Red circles* depict the RT of erroneous reaches directed first toward the emerging stimulus on Anti-reach trials. The EVR is the muscle recruitment falling between the *vertical white solid lines*, aligned to the point at which muscle recruitment differs depending on the side of stimulus emergence. The EVR is also depicted in the two rightmost columns as the activity falling within the *shaded interval* between the *vertical black solid lines*. All EMG activity is normalized to the level of activity over 500 ms preceding stimulus emergence. Note how EVRs increase following left stimulus emergence on Anti-reach trials, even though the reach proceeds to the right, and how EVRs decrease in the converse scenario.

Anti-reach trials dissociate the side of stimulus emergence from the direction of the ensuing reach. As reported previously (Gu et al., 2016; Kozak et al., 2020), the EVR on Anti-reach trials in our exemplar subjects was tied to the side of stimulus emergence, with EMG activity increasing following left stimulus emergence even when subjects correctly reached to the right and vice versa (third, fourth, and sixth columns in **Fig. 4**). The exemplar subjects occasionally reached incorrectly in the wrong direction on Anti-reach trials (trials with red dots in third and fourth columns), and when generated such movements tended to start earlier than correct reaches.

In both experiments, the prevalence of EVRs did not differ between PD patients and HCs (**Fig. 5A**). In Experiment 1, with at least 1500 ms of instruction time, all 16 PD patients and all 18 HCs generated an EVR in the Pro-reach condition. In Experiment 2 with 1000 ms of instruction time all 17 PD patients and 17 out of 18 HCs generated an EVR in the Pro-reach condition. In Experiment 2 with 500 ms of instruction time 16 out of 17 PD patients and 16 out of 18 HCs generated an EVR in the Pro-reach condition. A Chi-Square test of independence determined the prevalence of EVRs did not differ significantly between groups in Experiment 2 with 1000 ms (*X*^2^ (1, *N* = 35) = 0.972, *p* = 0.324) or 500 ms of instruction time (*X*^2^ (1, *N* = 35) = 0.305, *p* = 0.581). Furthermore, there was no effect of Group on EVR Latency following stimulus onset (**Fig. 5B**; *β* = 0.293, *SE* = 1.911, *p* = 0.879, *95% CI* [-3.453, 4.039]; *M_PD_* = 95.16 ms, *SD_PD_* = 8.547; *M_HC_* = 94.88 ms, *SD_HC_* = 8.277). The Bayes factor for the effect of Group indicated moderate support for the null hypothesis (*BF_10_* = 0.214). There was also no effect of Instruction time on EVR Latency (*β* = 0.443, *SE* = 1.494, *p* = 0.768, *95% CI* [-2.485, 3.370]; *M_1500_*= 93.53, *SD_1500_* = 6.947; *M_1000_* = 96.47, *SD_1000_* = 8.050; *M_500_* = 95.06 ms, *SD_500_* = 9.929). Together, these results indicate that EVRs are spared in patients with PD.

**Figure 5.**
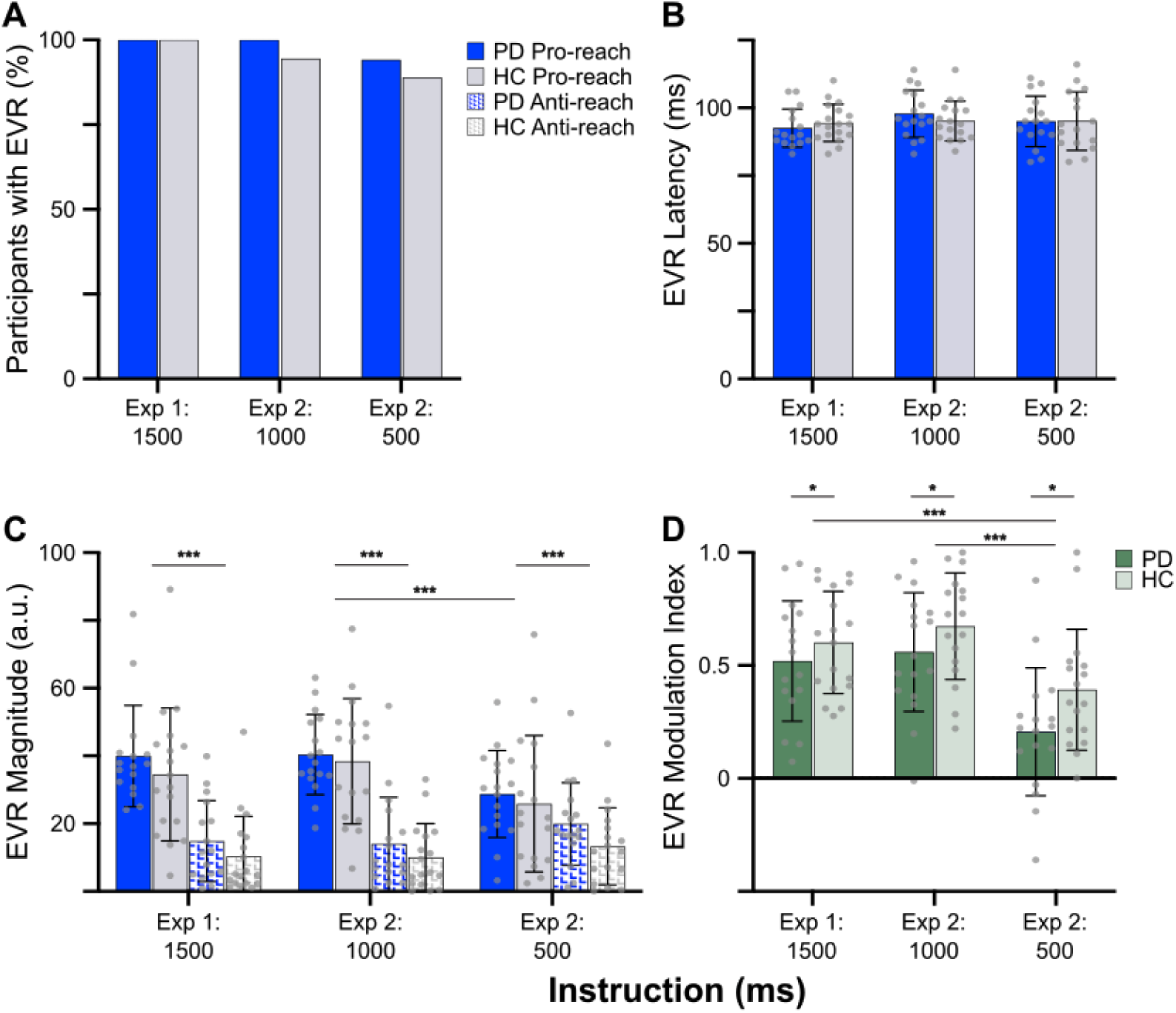
A. Patients with PD generated EVRs at the same prevalence as HCs with 1500 ms, 1000 ms, and 500 ms of instruction time. **B.** Patients with PD generated EVRs of the same latency as HCs with 1500 ms, 1000 ms, and 500 ms of instruction time. **C.** Participants had larger EVR Magnitudes on Pro-reach trials than Anti-reach trials. Participants had larger EVR Magnitudes with 1000 ms of instruction time than 500 ms of instruction time on Pro-reach trials. **D.** Patients with PD had lower EVR Modulation indices than HCs. Participants had greater EVR Modulation indices with 1500 ms and 1000 ms of instruction time than with 500 ms of instruction time. Bars represent *M*, except panel **A** in which they represent a percentage. Error bars represent *SD*, *\*p < 0.05*, *\*\*p < 0.01*, *\*\*\*p < 0.001*.

There was a significant main effect of Trial type on EVR Magnitude (**Fig. 5C**; *β* = 20.92, *SE* = 1.186, *p* < 0.001, *95% CI* [18.60, 23.24]), as participants generated larger EVRs on Pro-reach trials than on Anti-reach trials (*M_Pro_* = 34.57, *SD_Pro_*= 17.20; *M_Anti_* = 13.66, *SD_Anti_* = 12.08). There was no main effect of Instruction time (*β* = -3.365, *SE* = 1.914, *p* = 0.081, *95% CI* [-7.117, 0.387]; *M_1500_* = 24.75, *SD_1500_* = 19.39; *M_1000_* = 25.67, *SD_1000_*= 19.42; *M_500_* = 21.92, *SD_500_* = 15.43), however a significant interaction between Trial type and Instruction time (*β* = -15.35, *SE* = 2.515, *p* < 0.001, *95% CI* [-20.28, -10.42]) revealed that participants had larger EVRs with longer instruction on Pro-reach trials, but smaller EVRs with longer instruction on Anti-reach trials. This effect of Instruction time was significant for Pro-reach trials when comparing trials with 1000 ms of instruction time to those with 500 ms of instruction time (*p_Pro_* < 0.001; *p_Anti_* = 0.402), with no difference between 1500 ms and 1000 ms (*p_Pro_* = 1.000; *p_Anti_* = 1.000) or 1500 ms and 500 ms (*p_Pro_* = 0.088; *p_Anti_* = 1.000). There was no effect of Group on EVR magnitude (*β* = 4.225, *SE* = 3.264, *p* = 0.200, *95% CI* [-2.172, 10.62]). PD patients generated EVRs of similar magnitude compared to HCs (*M_PD_* = 26.25, *SD_PD_* = 16.72; *M_HC_* = 22.06, *SD_HC_* = 19.29). Although this did not reach significance in the linear mixed model, the Bayes factor for the effect of Group indicated some support for the alternative hypothesis, that patients with PD had larger EVRs than HC (*BF_10_*= 1.040).

### EVR Modulation

The Modulation index indicates the degree to which the task instruction influenced the magnitude of EMG activity during the EVR interval; a value of 1 or 0 indicates that subjects were completely able to or completely unable to suppress the EVR based on the Anti-reach instruction, respectively **(Fig. 5D)**. Patients with PD had significantly lower Modulation indices than HCs (*β* = -0.127, *SE* = 0.059, *p* = 0.035, *95% CI* [-0.243, -0.011]; *M_PD_* = 0.427, *SD_PD_* = 0.310; *M_HC_* = 0.557, *SD_HC_* = 0.267), indicating an impairment in the ability of task instruction to influence the EVR. There was also a significant main effect of Instruction time on the Modulation index (*β* = -0.289, *SE* = 0.043, *p* < 0.001, *95% CI* [-0.374, -0.205]). As expected, participants had more EVR modulation with 1500 ms (*p* < 0.001) and 1000 ms (*p* < 0.001) compared to only 500 ms of instruction time, and there was no difference between 1500 ms and 1000 ms (*p* = 1.000; *M_1500_* = 0.563, *SD_1500_* = 0.245; *M_1000_* = 0.617, *SD_1000_* = 0.252; *M_500_* = 0.300 ms, *SD_500_* = 0.287).

### Muscle recruitment after the EVR: Reach burst magnitude

The EVR is the first wave of muscle recruitment, influenced by stimulus emergence. As shown in the left column of **Fig. 6A** for data aligned to stimulus onset, muscle recruitment persists for some time after the EVR through movement initiation (see also **Fig. 4**). To examine muscle recruitment prior to reach onset, we realigned all data to reach onset (right column in **Fig. 6A**), and then derived the Reach burst magnitude as the integral of EMG activity over 30 ms centered on the peak difference for left vs right reaches. Two outliers in the HC group were removed from this analysis which were identified as participants with values more than 3 *SD* above the group mean. The effect of Group on Reach burst magnitude was significant (**Fig. 6B**; *β* = -17.75, *SE* = 7.404, *p* = 0.020, *95% CI* [-32.26, -3.241]). PD patients had less EMG activity in the interval immediately preceding movement onset (*M_PD_* = 82.09, *SD_PD_* = 23.89; *M_HC_* = 99.79, *SD_HC_* = 33.46). There was no main effect of Trial type, as participants had similar EMG activity preceding pro- and anti-reaches (*β* = -0.264, *SE* = 1.659, *p* = 0.874, *95% CI* [-3.515, 2.987]; *M_Pro_* = 90.98, *SD_Pro_* = 28.43; *M_Anti_* = 91.16, *SD_Anti_* = 32.33). There was also no effect of Instruction time on Reach burst magnitude. Participants had similarly sized reach bursts with 1500 ms, 1000 ms, and 500 ms of instruction time (*β* = 2.837, *SE* = 3.860, *p* = 0.464, *95% CI* [-4.729, 10.40]; *M_1500_* = 90.09, *SD_1500_* = 33.70; *M_1000_* = 90.16, *SD_1000_* = 28.28; *M_500_* = 92.80, *SD_500_* = 29.33).

**Figure 6.**
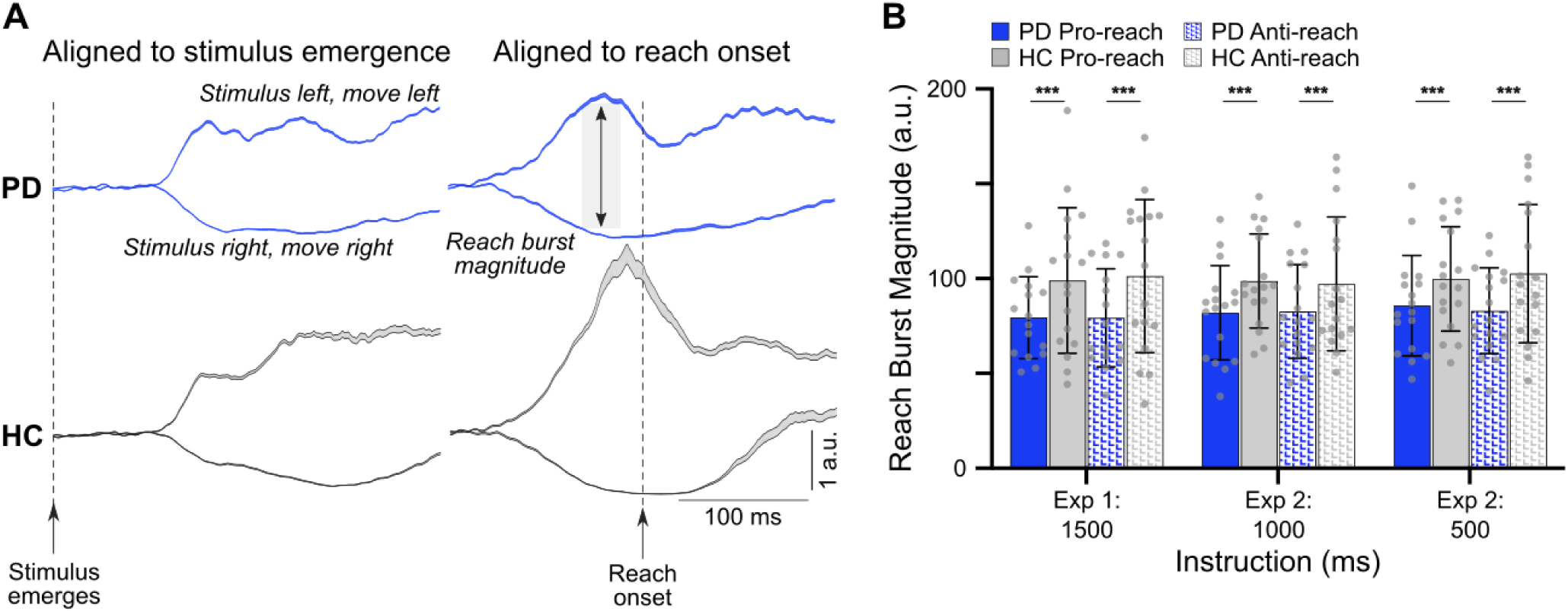
Calculation of recruitment magnitude aligned to reach onset is shown for PD and HCs. **A.** Contours depict average EMG recruitment for pro-reaches across all PD and HC participants, aligned to either stimulus emergence (left column) or reach onset (right column). All data were normalized to recruitment in the 100 ms preceding stimulus emergence. Reach burst magnitude is calculated as the integral over 30 ms, centered on the peak difference between activity for left vs. right reaches (*light grey rectangle*) preceding reach onset. **B.** Reach burst magnitudes. Patients with PD had smaller Reach burst magnitudes than HCs. Bars represent *M*, error bars represent *SD*, *\*p < 0.05*, *\*\*p < 0.01*, *\*\*\*p < 0.001*.

### Correlations of EVR and Reach Burst Magnitude with Behavioural Measures of Reaching

Finally, we were interested in whether there were correlates of muscle recruitment during the EVR interval and the reach burst interval with behavioural measures (**Table 2**). In PD, larger mean amplitude EVRs on Pro-reach trials were negatively correlated with RT at 1500 ms (*r*(14) = -0.670, *p* = 0.004), and 500 ms (*r*(15) = -0.801, *p* < 0.001). Similarly, in HC larger EVRs on Pro-reach trials were strongly negatively correlated with RT at 1500 ms (*r*(14) = -0.709, *p* < 0.001), 1000 ms (*r*(15) = -0.884, *p* < 0.001), and 500 ms (*r*(15) = -0.875, *p* < 0.001). The Modulation index of the EVR was strongly a) negatively correlated with Error rate in the Anti-reach condition and b) positively correlated with *d’*, our measure of response regulation based on Pro- versus Anti-reach instruction. In PD, these correlations reached significance at 1500 ms [*r*(14) = -0.703, *p* = 0.002 and *r*(14) = 0.654, *p* = 0.006, respectively], 1000 ms [*r*(15) = -0.841, *p* < 0.001 and *r*(15) = 0.816, *p* < 0.001, respectively], and 500 ms [*r*(15) = -0.772, *p* < 0.001 and *r*(15) = 0.851, *p* < 0.001, respectively]. In HC, the Modulation index correlated with negatively with Error rate in the Anti- reach condition and positively with *d’* at 1000 ms [*r*(16) = -0.648, *p* = 0.005 and *r*(16) = 0.617, *p* = 0.008, respectively], and positively with Peak velocity and negatively with Movement duration in the Pro-reach condition at 500 ms [*r*(16) = 0.659, *p* = 0.004 and *r*(16) = -0.620, *p* = 0.008, respectively].

**Table 2:** Correlations of muscle recruitment with behavioural measures of reaching

|  |  | 1500 ms |  |  | 1000 ms |  |  | 500 ms |  |  |
| --- | --- | --- | --- | --- | --- | --- | --- | --- | --- | --- |
|  |  | EVR Magnitude (Pro-Reach) | Modulation Index | Reach Burst Magnitude (Pro-reach) | EVR Magnitude (Pro-Reach) | Modulation Index | Reach Burst Magnitude (Pro-reach) | EVR Magnitude (Pro-reach) | Modulation Index | Reach Burst Magnitude (Pro-reach) |
| PD | RT (Pro-reach) | **-0.670**<br>$p = 0.004$ | -0.050<br>$p = 0.855$ | -0.123<br>$p = 0.651$ | -0.572<br>$p = 0.016$ | 0.045<br>$p = 0.865$ | -0.256<br>$p = 0.321$ | **-0.801**<br>$p < 0.001$ | -0.353<br>$p = 0.165$ | -0.049<br>$p = 0.851$ |
| | Error Rate (Anti-reach) | 0.094<br>$p = 0.729$ | **-0.703**<br>$p = 0.002$ | -0.030<br>$p = 0.912$ | -0.022<br>$p = 0.935$ | **-0.841**<br>$p < 0.001$ | -0.406<br>$p = 0.106$ | 0.000<br>$p = 0.999$ | **-0.772**<br>$p < 0.001$ | -0.554<br>$p = 0.021$ |
| | C | -0.340<br>$p = 0.197$ | 0.575<br>$p = 0.020$ | -0.036<br>$p = 0.895$ | -0.044<br>$p = 0.867$ | 0.347<br>$p = 0.173$ | 0.139<br>$p = 0.594$ | -0.535<br>$p = 0.027$ | 0.166<br>$p = 0.525$ | 0.545<br>$p = 0.024$ |
| | $d'$ | 0.004<br>$p = 0.989$ | **0.654**<br>$p = 0.006$ | -0.038<br>$p = 0.890$ | 0.039<br>$p = 0.881$ | **0.816**<br>$p < 0.001$ | 0.406<br>$p = 0.106$ | 0.236<br>$p = 0.362$ | **0.851**<br>$p < 0.001$ | 0.410<br>$p = 0.102$ |
| | Peak Velocity (Pro-reach) | -0.314<br>$p = 0.236$ | 0.072<br>$p = 0.792$ | 0.383<br>$p = 0.143$ | -0.026<br>$p = 0.921$ | 0.453<br>$p = 0.068$ | 0.440<br>$p = 0.077$ | -0.067<br>$p = 0.798$ | 0.485<br>$p = 0.048$ | 0.310<br>$p = 0.226$ |
| | Movement Duration (Pro-reach) | 0.278<br>$p = 0.298$ | 0.100<br>$p = 0.712$ | -0.307<br>$p = 0.248$ | 0.179<br>$p = 0.491$ | -0.417<br>$p = 0.096$ | -0.343<br>$p = 0.177$ | 0.186<br>$p = 0.474$ | -0.507<br>$p = 0.038$ | -0.305<br>$p = 0.233$ |
| HC | RT (Pro-reach) | **-0.709**<br>$p < 0.001$ | -0.154<br>$p = 0.542$ | 0.352<br>$p = 0.152$ | **-0.884**<br>$p < 0.001$ | 0.552<br>$p = 0.022$ | 0.359<br>$p = 0.157$ | **-0.875**<br>$p < 0.001$ | 0.225<br>$p = 0.384$ | 0.046<br>$p = 0.861$ |
| | Error Rate (Anti-reach) | 0.417<br>$p = 0.085$ | -0.302<br>$p = 0.224$ | -0.114<br>$p = 0.653$ | 0.609<br>$p = 0.010$ | **-0.648**<br>$p = 0.005$ | 0.275<br>$p = 0.286$ | 0.615<br>$p = 0.009$ | -0.443<br>$p = 0.075$ | -0.117<br>$p = 0.655$ |
| | C | **-0.386**<br>$p = 0.114$ | 0.155<br>$p = 0.539$ | -0.352<br>$p = 0.512$ | -0.343<br>$p = 0.178$ | 0.434<br>$p = 0.082$ | -0.407<br>$p = 0.105$ | -0.278<br>$p = 0.280$ | 0.393<br>$p = 0.119$ | 0.326<br>$p = 0.202$ |
| | $d'$ | -0.344<br>$p = 0.162$ | 0.353<br>$p = 0.151$ | -0.037<br>$p = 0.886$ | **-0.620**<br>$p = 0.008$ | **0.617**<br>$p = 0.008$ | -0.293<br>$p = 0.254$ | **-0.710**<br>$p = 0.001$ | 0.528<br>$p = 0.029$ | 0.151<br>$p = 0.562$ |
| | Peak Velocity (Pro-reach) | -0.246<br>$p = 0.325$ | -0.240<br>$p = 0.337$ | **0.880**<br>$p < 0.001$ | -0.544<br>$p = 0.024$ | 0.268<br>$p = 0.298$ | **0.709**<br>$p < 0.001$ | -0.462<br>$p = 0.062$ | **0.659**<br>$p = 0.004$ | **0.777**<br>$p < 0.001$ |
| | Movement Duration (Pro-reach) | 0.033<br>$p = 0.897$ | 0.019<br>$p = 0.940$ | **-0.641**<br>$p = 0.004$ | 0.468<br>$p = 0.058$ | -0.312<br>$p = 0.223$ | **-0.666**<br>$p = 0.004$ | 0.430<br>$p = 0.085$ | **-0.620**<br>$p = 0.008$ | **-0.706**<br>$p = 0.002$ |

**Note:** Data are presented as Pearson correlation coefficients ( $r$ ) and $p$ values; $*p < .008$ , alpha value Bonferroni-corrected.

In contrast, the second phase of the EMG activity, the Reach burst magnitude, was not correlated with RT, Anti-Reach Error rate, *C*, or *d’*. Rather, in HC, the Reach burst magnitude was strongly a) positively correlated with Peak velocity and b) negatively correlated with Movement duration at all instruction intervals [*r*(16) = 0.880, *p* < 0.001 and *r*(16) = -0.641, *p* = 0.005 at 1500 ms; *r*(16) = 0.709, *p* < 0.001 and *r*(16) = -0.666, *p* = 0.004 at 1000 ms; *r*(16) = 0.777, *p* < 0.001 and *r*(16) = -0.706, *p* = 0.002 at 500 ms, respectively]. This reveals a double dissociation whereby the EVR impacts measures of reach initiation such as RT, Error rate, C, and *d’,* and the second burst of EMG activity, the Reach burst magnitude, relates to the measures of reach execution speed such as the Peak velocity and Movement duration. These correlations with Reach burst magnitude did not reach significance in PD, likely due to increased variability in Reach burst magnitude in patients.

## Discussion

PD patients (mean disease duration: Experiment 1 = 3.80 years +/- 4.68, Experiment 2 = 5.87 years +/- 4.62; mean MDS-UPDRS: Experiment 1 = 31.69 +/- 9.79, Experiment 2: 28.53 +/- 13.55; mean LEDD: Experiment 1 = 479.69 mg +/- 354.17, Experiment 2 = 633.41 mg +/- 332.13) abstaining from their usual dopaminergic therapy, and age-matched HCs performed visually-guided reaches either toward or away from a stimulus in the Emerging target paradigm. In Experiment 1, the interval between Pro-reach or Anti-reach instructions and stimulus emergence was at least 1500 ms. In Experiment 2, we increased task difficulty and time pressure to adjust to Pro- or Anti-reach instructions, reducing the instruction-to-stimulus emergence interval to 1000 ms or 500 ms. In both experiments, we observed two distinct phases of muscle recruitment in the upper limb that related well to different aspects of our kinematic and behavioural results, some (but not all) of which were spared in PD. The first phase of muscle recruitment—the stimulus-evoked EVR—was remarkably similar in magnitude and latency in PD and HC; sparing of the EVR related well to the similar reaction times in PD and HC. However, PD patients exhibited less contextual suppression of the EVR on Pro- vs Anti-reach trials, and the impaired contextual suppression correlated with the increased Anti-reach Error rate and lower response regulation (i.e., *d’*) in PD. After the EVR, a second and longer-lasting phase of muscle recruitment persisted up to and beyond reach initiation. When aligned to reach onset, this second phase of recruitment was greatly reduced in PD; PD patients exhibited lower Peak velocities and longer Movement duration. Below, we expand on the likely descending motor circuits engaged in this task and the impact of PD on such circuits, and the implications of our results on the understanding of KP and the bradykinetic presentation of PD.

### Insights gained from a trial-by-trial analysis of limb muscle recruitment

The insights of this study are predicated on trial-by-trial recording of EMG signals from an upper limb muscle, which in the posture and movement studied here, unfolds over hundreds of milliseconds. EMG measures provide direct insights into neuromuscular control of reach initiation and execution, and indirect insights into movement preparation by virtue of assessing how these signals change on a particular type of trial. The behavioural paradigm incorporates aspects of both bottom-up (e.g., stimulus emergence) and top-down (e.g., preparing for a Pro- or Anti-Reach with various instruction intervals) control, and by assessing such signals in both PD patients and HC, we can appreciate the selective nature of PD pathophysiology on the nested descending motor control circuits engaged in this visually-guided reaching task.

The EVR precludes substantive processing of visual stimuli through cortical circuits, and previous work has built a circumstantial case that EVR provides a unique window into signalling along the tecto-reticulospinal pathway (Corneil et al., 2004; Boehnke and Munoz, 2008; Pruszynski et al., 2010; Corneil and Munoz, 2014), preceding the volleys of recruitment arising through descending circuits originating in the cortex. Previous investigations conducted exclusively in a younger cohort has emphasized that this pathway can be contextually pre-set by top-down control, influencing EVR magnitude but not latency (Gu et al., 2016; Contemori et al., 2023). Our current study is the first conducted in an elderly cohort, and all PD patients and all but one HC generated EVRs. All aspects of the trial-by-trial variation in the EVR on Pro- and Anti-Reach trials, and its relationship to RTs and Anti-Reach errors, replicated previous reports from a younger cohort (Gu et al., 2016; Kozak et al., 2020). Our current findings emphasize that the pathway mediating the EVR, which appears to arise from the shortest pathway by which visual inputs can influence reaching actions (Gu et al., 2018; Weerdesteyn et al., 2024), is functional in an elderly cohort and spared in PD.

Although EVR Latency and Magnitude were unaffected by PD, PD patients exhibited less ability to contextually suppress the EVR on Anti-reach trials. Thus, the brain circuits generating the EVR are dissociable from those regulating the EVR, with the latter circuits being selectively impacted by the nigro-striato-cortical pathophysiology of PD. The movement-related layers of the SC from which the tecto-reticulospinal tract takes origin receives convergent top-down inputs from numerous fronto-parietal areas (May, 2006; Gandhi and Katnani, 2011), with such cortical inputs influencing the contextual expression but not production of reflexive movements like EVRs and express saccades (Schiller et al., 1987; Munoz et al., 2000; Rezvani and Corneil, 2008; Chen et al., 2013; Dash et al., 2018). The behavioural consequence of the EVR, and the importance of proper regulation, are apparent in our kinematic results. The RTs of movement toward a salient visual stimulus (i.e., Pro-reaches) are spared in PD but such stimulus-driven movements are incongruent with task instruction on Anti-reach trials, and the inability to appropriately suppress the EVR is apparent in the increased Error rate in PD on Anti-reach trials. Across the PD and HC sample, we observed considerable variability in the Modulation index (Fig. 5D), and the strong correlations of this value to measures like Anti-reach Error rate and response regulation (*d’*) reinforce the behavioural relevance of forces arising from the EVR.

While important for response initiation, EVR Magnitude did not correlate with other kinematic measures like Peak velocity or Movement duration, which as expected, were strongly compromised in PD. Instead, the bradykinetic presentation in PD is related to a decreased Reach burst magnitude, which measures the vigor of muscle recruitment around the time of reach onset. Given the nested and parallel nature of descending motor control circuits in primates (Lemon, 2008; Alstermark and Isa, 2012), the bradykinetic presentation in PD likely arises from deficient corticospinal inputs directly onto motoneurons, and indirectly from deficient cortical inputs to spinal and supraspinal networks, including in the reticular formation (Tapia et al., 2022), which then drive the motoneurons. The overall presentation in PD of spared movement initiation but compromised movement velocities resembles that reported recently in an interceptive task (Fooken et al., 2022) for PD patients on dopaminergic medication. The PD cohort we studied refrained from their usual dopaminergic therapy prior to data collection, and a future study will detail the impact of dopaminergic medication on stimulus-aligned and movement-aligned phases of upper limb muscle recruitment.

### Implications of our results for the understanding of Kinesis Paradoxica (KP)

Our results speak to the phenomenon of KP, wherein swift and fluid movements can be elicited even in advanced PD patients by suddenly-presented and salient stimuli, by physically threatening contexts, or by engaging well-rehearsed motor routines such as cycling or skiing (Daroff, 2008; Snijders et al., 2011; Nonnekes et al., 2019; Duysens and Nonnekes, 2021; Melo-Thomas and Schwarting, 2023; Tostes et al., 2024). Visual, auditory, or tactile stimuli can also help interrupt episodes of freezing, induce gait initiation, and improve gait speed and quality (Dunne et al., 1987; Thaut et al., 1996; Suteerawattananon et al., 2004; Jiang and Norman, 2006; Bieńkiewicz et al., 2013, 2014; Cassimatis et al., 2016; An et al., 2023). In experimental settings, KP has been most commonly studied using conspicuous (e.g., abruptly onsetting, moving, high-contrast) stimuli, resembling the visual stimuli we used. For example, PD patients can rapidly grasp a suddenly-appearing ball that rolls down a ramp, despite impairments in generating such movements in response to a stationary ball (Majsak et al., 1998). PD patients can also rapidly and accurately adjust on-going reaching movements in response to a jumped target (Merritt et al., 2017), or reach to intercept moving stimuli (Fooken et al., 2022).

Recognized 100 years ago, the mechanism(s) of KP are still not clearly established (Duysens and Nonnekes, 2021; Fasano et al., 2022; Melo-Thomas and Schwarting, 2023). Our findings challenge the long-held proposition that KP arises from switching away from habitual toward voluntary motor control through increased concentration or attention. Doing so is proposed to normalize movement by breaking out of and by-passing striatal circuits impaired in PD that purportedly mediate habitual actions and automatic motor programs (Glickstein and Stein, 1991; Morris, 2000; Redgrave et al., 2010; Wu et al., 2015; Hess and Hallett, 2017; Nonnekes et al., 2019; Bologna et al., 2020; Fasano et al., 2022; Melo-Thomas and Schwarting, 2023). This explanation appears largely inconsistent with our core observations of spared EVRs but impaired regulation of the EVR on Anti-reach trials, and deficits in subsequent volleys of recruitment following the EVR.

Another explanation for KP proposes an important role for the pontine nuclei (Glickstein and Stein, 1991; Fasano et al., 2022; Melo-Thomas and Schwarting, 2023), given the responsiveness of these nuclei to stimuli that provoke KP, and the circuitry that bypasses the dopamine-depleted striatum through the cerebellum and motor cortex (Baker et al., 1976; Suzuki and Keller, 1984; Suzuki et al., 1990; Karbasforoushan et al., 2022). This explanation seems inconsistent with recent results in rodents showing that lesions of the pontine nuclei affected reaching kinematics without disrupting initiation latency (Guo et al., 2021), and is also inconsistent with both the rapid and unchanged latency of the EVRs in PD reported here, and the fact that it is the subsequent phases of recruitment after the EVR, which presumably do involve signalling through motor cortex, that are impaired.

Instead, our results are compatible with hyper-habitual responding and a failure to invoke more controlled and flexible responses in PD, rather than adoption of voluntary cortical motor pathways to compensate for impairment in overlearned stimulus-response behaviours mediated by striatal motor circuits. In the context of our experiment, we suggest that a fast, subcortical motor pathway mediates KP by producing reflexive reaches toward an emerging visual stimuli, even when participants are instructed to move in the opposite direction. The visual stimuli known to preferentially evoke the EVR also robustly generate visual responses in the movement-related layers of the superior colliculus (Wood et al., 2015; Kozak et al., 2019; Kozak and Corneil, 2021). The SC has long been implicated in oculomotor control, but we and others have argued that EVRs on the neck (Corneil et al., 2004), upper limb (Pruszynski et al., 2010; Contemori et al., 2021a), and lower (Fautrelle et al., 2010; Billen et al., 2023) limb are the skeletomotor equivalents of express saccades. Indeed, there are striking similarities between the properties of the upper limb muscle recruitment we observed in PD patients and oculomotor responses in a variety of eye movement tasks, which have broadly observed a sparing of express saccades, largely spared RTs for pro-saccades, an increased propensity for saccade errors in the Anti-saccade task, and difficulty generating internally-generated eye movements (Briand et al., 1999; Chan et al., 2005; Terao et al., 2011, 2013; Pretegiani and Optican, 2017; Waldthaler et al., 2021; Fooken et al., 2022; Riek et al., 2023; Antoniades and Spering, 2024). Unlike ballistic eye movements, close assessment of EMG activity allows resolution of stimulus-driven and internally-generated volleys of recruitment within a single trial.

The inferior colliculus (IC) is also commonly invoked as mediating KP (Melo-Thomas and Schwarting, 2023), given its strong auditory inputs and the well-known benefits of music and rhythmic auditory cueing for patients suffering from freezing of gait (Nieuwboer et al., 2009; Rochester et al., 2009; Nombela et al., 2013). The IC is also connected to the medial descending motor systems, and processing through the IC or SC may be relevant for auditory or visual stimuli respectively in KP. Further, the movement-related layers of the SC are highly multimodal (Stein and Stanford, 2008), and reticulospinal outputs can be triggered with visual, auditory, vestibular, or tactile inputs (Glover and Baker, 2019; Weerdesteyn et al., 2024). Our findings of spared EVRs resonate with other work showing that startle mediated phenomena thought to depend on the reticular formation are also spared in PD (Valldeoriola et al., 1998; Nonnekes et al., 2015). Such spared subcortical circuits would appear to provide the requisite connection to the medial descending motor systems to mediate KP. Indeed, such subcortical circuits, and the SC itself, may also be engaged by threatening or emotional stimuli, given that KP can be provoked in emotionally charged situations like earthquakes (Bonanni et al., 2010), fires (Jankovic, 2008), or nightmares (Tostes et al., 2024). Explanations from these case studies suggest that such situations trigger an emotional state that increases attention, focus, or internal motivation to move. Alternatively, such situations may trigger strong activation of the innate alarm system (Olivé et al., 2018), and initiate movement through the medial descending motor systems via connections through the locus coeruleus to the superior colliculus and/or reticular formation.

KP can be elicited in a variety of scenarios. Such observations have led to explanations that invoke processing through cortical areas. Instead, our results show that an evolutionary-conserved subcortical circuit is intact in PD which, when activated by stimuli that provoke KP, can produce normal movement. The integrity of this circuit, and the converging inputs it receives from multisensory and affective areas, offers an alternative perspective for movement initiation in KP. Once initiated, and depending on the type of movements being generated, the movements themselves may trigger abundant sources of sensory reafference that help sustain movement, at least for some period of time.

## GRANTS

This work was supported by a seed grant to BDC and PAM from the Schulich School of Medicine and Dentistry, as well as funds from the Canadian Institutes of Health Research (CIHR; PAM: R4981A15; BDC: MOP-93796, -142317; PJT-180279), Natural Sciences and Engineering Research Council of Canada (BDC RGPIN:-311680, -04394-2021), Canada Research Chairs program (PAM: 950-230372), and equipment funds from the Canadian Foundation for Innovation. RAK and MG were supported by an Ontario Graduate Scholarship, and MG was supported by a CIHR Canada Graduate Scholarship, and funding from the Parkinson’s Society of Southwestern Ontario.

## Notes

### Competing Interest Statement

The authors have declared no competing interest.

